# Evolution of hierarchy and irreversibility in theoretical cell differentiation model

**DOI:** 10.1101/2023.07.04.547517

**Authors:** Yoshiyuki T. Nakamura, Yusuke Himeoka, Nen Saito, Chikara Furusawa

## Abstract

The process of cell differentiation in multicellular organisms is characterized by hierarchy and irreversibility in many cases. However, the conditions and selection pressures that give rise to these characteristics remain poorly understood. By using a mathematical model, here we show that the network of differentiation potency (differentiation diagram) becomes necessarily hierarchical and irreversible by increasing the number of terminally differentiated states under certain conditions. The mechanisms generating these characteristics are clarified using geometry in the cell state space. The results demonstrate that the appearance of these characteristics can be driven without assuming the adaptive significance. The study also provides a new perspective on the structure of gene regulatory networks that produce hierarchical and irreversible differentiation diagrams. These results indicate some constraints on cell differentiation, which are expected to provide a starting point for theoretical discussion of the implicit limits and directions of evolution in multicellular organisms.

## INTRODUCTION

Multicellular organisms have various cell types that are generated during the process of cell differentiation. The process of cell differentiation can be characterized by the following important properties, namely hierarchy and irreversibility [1]. Typically, this process involves a series of hierarchical fate decisions before reaching a final non-differentiable state (“hierarchy”). Once a cell reaches its downstream state, it cannot revert back to its stem cell state during normal development (“irreversibility”). These features of cell differentiation are commonly seen in various species [2–10]. Waddington’s “epigenetic landscape” is a qualitative metaphor that captures these characteristics of the cell differentiation process [11]. Understanding the cell state transitions described above remains a key goal in developmental biology, and has also recently become a significant challenge in the context of medical applications, such as reprogramming [12] and cancer therapy [13, 14].

To theoretically understand cell differentiation, mathematical models have been employed and developed. These models are usually based on the fact that the gene expression state of a cell is regulated by a gene regulatory network (GRN) [15–17]. For instance, Kauffman linked cell types to multiple attractors in a dynamical system of a Boolean network model that abstracts a GRN [18, 19]. In this model, the expression state of each gene is represented as a binary value and the cell state is defined as a vector composed of these values. Despite its bold formulation, the model has been successful in explaining gene expression data in real organisms [20, 21]. Based on this pioneering work, further theoretical models have been established that consider transitions between cell types. For example, Huang and colleagues demonstrated that bistable switches, composed of two self-activating and mutually repressing genes, can theoretically exhibit a differentiation process into two distinct differentiated types from an identical stem cell type [22, 23].

Moreover, several models have been proposed that can generate hierarchical and irreversible networks of differentiation potency (differentiation diagrams) between multiple stable states. For instance, hierarchical differentiation diagrams can be constructed when two or more bistable switches are connected in a hierarchical manner [23–27]. Other mechanisms, such as Turing instability [28, 29], temporal change of an expression system assuming epigenetic changes [30] and bifurcation of dynamical systems [31] have also been proposed to design hierarchical and irreversible differentiation diagrams.

As demonstrated thus far, mechanisms capable of exhibiting hierarchy and irreversibility have been studied and proposed, but the reasons why these characteristics are commonly observed among various species remain poorly understood. Specifically, what conditions and selection pressures give rise to hierarchy and irreversibility? By what mechanisms are they acquired? As differentiation diagrams with hierarchy and irreversibility are the outcome of evolution from simpler structures, it is required to investigate how these diagrams change their topology during their evolution. In order to answer these questions, a theoretical study is necessary that is independent of specific species details.

In view of taking cell types as attractors of dynamical systems, the structure of the differentiation diagram can be reframed as a question of how the basin boundaries of attractors are arranged. If it is assumed that transitions between them can occur due to some kind of perturbation (e.g., fluctuations in gene expression, signal-induced changes in expression levels or gene-gene interaction), understanding the arrangement of the basin boundaries can help to understand which basins the cell state can move between.

Here we use an abstract cell model with a GRN to investigate how hierarchy and irreversibility emerge in cell differentiation during evolution. In our study, the multicellular systems are optimized to increase the number of terminally differentiated states, envisioning an abstract evolutionary process. We vary the magnitude of perturbations to gene expression driving differentiation and show that large perturbation conditions lead to inevitable hierarchical differentiation, whereas small perturbation conditions do not. Furthermore, under the conditions that lead to the emergence of a hierarchical differentiation diagram, the diagrams also become irreversible. These findings indicate that hierarchy and irreversibility are inherent outcomes of the evolutionary process under certain conditions. We also clarify the mechanism for their inevitable emergence by analyzing the geometrical aspects of the cell state space. Finally, we explore the typical structures observed in GRNs resulting in differentiation diagrams with hierarchy and irreversibility.

Overall, our findings suggest the possibility that the nature of cell differentiation that emerged through evolution is due to secondary effects resulting from the optimization of the number of terminally differentiated states, rather than adaptive significance. In addition, our results offer a new perspective on the structure of GRNs that exhibit hierarchy and irreversibility.

## MODEL

To investigate the evolution of differentiation diagrams, we consider a cell model where cells have the capability to differentiate into another cell type (Fig. 1).

**FIG. 1.**
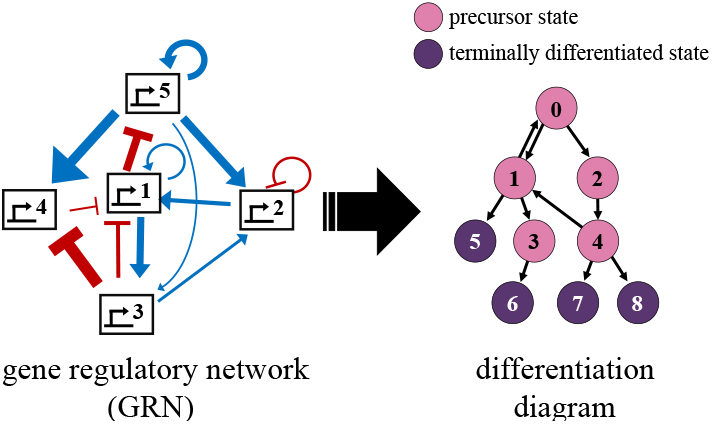
A schematic of this model. (left) An abstract representation of a GRN. The blue arrows represent positive interaction, while the red arrows represent negative interactions. (right) An abstract representation of the differentiation diagram. Each node represents an attractor of the dynamical system determined by the GRN on the left. The arrows represent possible differentiation paths by perturbation. The pink nodes represent precursor states, while the purple nodes represent terminally differentiated states.

### Cell state regulation

The model cell is described by a gene regulatory network (GRN) in which each gene expression is regulated by the other genes. The state of a cell, denoted by ***x*** = (*x*_1_, *x*_2_, … *x*_*n*_) is characterized by the expression levels of *n* genes. The dynamics of a cell state ***x*** obeys the following ordinary differential equation.

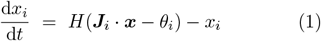

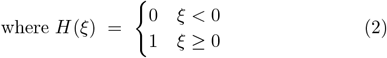

The matrix *J* represents the GRN, where its element *J*_*ij*_ represents how strongly gene *j* interacts with gene *i*. ***J***_*i*_ is the *i*-th row of *J* . *θ*_*i*_ is the expression threshold for each gene *i*. When the sum of the interactions from all genes (***J***_*i*_ ***x***) exceeds the threshold *θ*_*i*_, the first term *H* (·) takes 1, indicating that the mode of expression of gene *i* is “on”. The second term represents the decay of gene expression. Because of this term, the range of ***x*** is limited to [0,1].

### Differentiation by perturbation on expression

Fixed point attractors in the dynamical system according to eq. (1) are considered to represent distinct cell types. To determine the directions in which differentiation is possible between these cell types, we need to consider the mechanisms driving differentiation. While there are several possible driving forces that actually cause differentiation, we assume for simplicity that differentiation is caused by stochastic fluctuations in gene expression [23, 32]. We then derive a differentiation diagram using the following method.

The attractor to which an initial point ***x***^init^ = **0** converges is referred to as “the root attractor (***a***^root^)”. Throughout this research, we assume that ***a***^root^ represents the most upstream stem cell type (e.g., hematopoietic stem cell). Since the expression states of 0 and 1 are symmetric in this model and since we later evolve the differentiation diagram with ***x***^init^ fixed, choosing **0** for ***x***^init^ has no essential meaning.

Now, to define differentiation by perturbation, we consider adding a perturbation ***p***_*k*_ to an attractor ***a*** to create a new state ***a*** + ***p***_*k*_ for all *k* (= 1, 2, *N*), where *N* is the number of perturbations. If this new state subsequently converges to another attractor ***a***^*′*^ (≠ ***a***), we consider it possible to differentiate from ***a*** to ***a***^*′*^. Starting from ***a***^root^, we repeat the same procedure for all newly emerged attractors until no new attractor is found any further. Note that a differentiation diagram can be derived by determining *J* and ***θ***.

If no further transition occurs from a certain attractor under given perturbations, the attractor is referred to as the “terminally differentiated state” hereafter. This means that after any perturbation is added, the new initial state converges back to the attractor before the perturbation. Attractors that are not the terminally differentiated state are referred to as “precursor states”.

The subsequent results presented in this study are relatively independent of the number of perturbations (*N*), as long as it is sufficiently large. In this research, *N* = 1000 was used. Here, we assume that the norm of every perturbation vector, ||***p***_*k*_|| is identical, which is represented as a parameter *δ*. Thus, each ***p***_*k*_ is uniformly distributed on a hypersphere with radius *δ*. In this research, we will investigate the changes in the differentiation diagram under various conditions of *δ* while optimizing the number of terminally differentiated states using the method described below.

### Optimization of GRN

In this study, we aim to increase the number of terminally differentiated states by optimizing both the GRN matrix, denoted by *J*, and the threshold vector, denoted by ***θ***, using a genetic algorithm (GA). In other words, the fitness of the GA is set as the number of terminally differentiated states obtained. The reason for this is based on our assumption that the diversity of terminally differentiated states is linked to the number of specialized functions of cells. This is supported by the fact that many specialized cell types, such as neurons, osteoblasts, and muscle cells, do not undergo further differentiation [33]. Taking this into account, the above-mentioned fitness is set, guided by the idea that species with a higher number of terminally differentiated states, reflecting diverse functionalities, possess evolutionary advantages.

Furthermore, we also randomly sample high-fitness (*J*, ***θ***), using multicanonical sampling [34], a type of Markov chain Monte Carlo method. The reason we use this method is to collect a number of rare high-fitness samples efficiently. Using this method for a sufficiently long time, we can sample diagrams as a random walk along the fitness axis, without getting trapped in local optima. [34].

## RESULTS

### Acquisition of hierarchy

To investigate the influence of *δ* on the differentiation diagram, particularly on its hierarchical structure, we implement GA under various conditions of *δ*. Fig. 2a shows typical differentiation diagrams which evolved under two different *δ* values. Under *δ* = 0.05, almost all terminally differentiated states directly emerge from the root attractor, i.e., little hierarchy is observed in the differentiation diagram. In contrast, in the cases under *δ* = 0.4, the differentiation diagram generally becomes more complex and hierarchical. Notably, the final mean fitness does not exhibit a significant change when *δ* varies between 0.05 to 0.4 (Fig. 2b). However, it become much more difficult to increase the fitness when *δ ≥* 0.5 since the perturbations are too large. In the region where evolution is reasonably possible (here, with a final average fitness of around 15 or more, which is 0.05 ⪅ *δ* ⪅ 0.4), a larger *δ* leads to a greater value of depth (Fig. 2b). We will focus mainly on the results obtained under the conditions of *δ* = 0.05 and 0.4 hereafter.

**FIG. 2.**
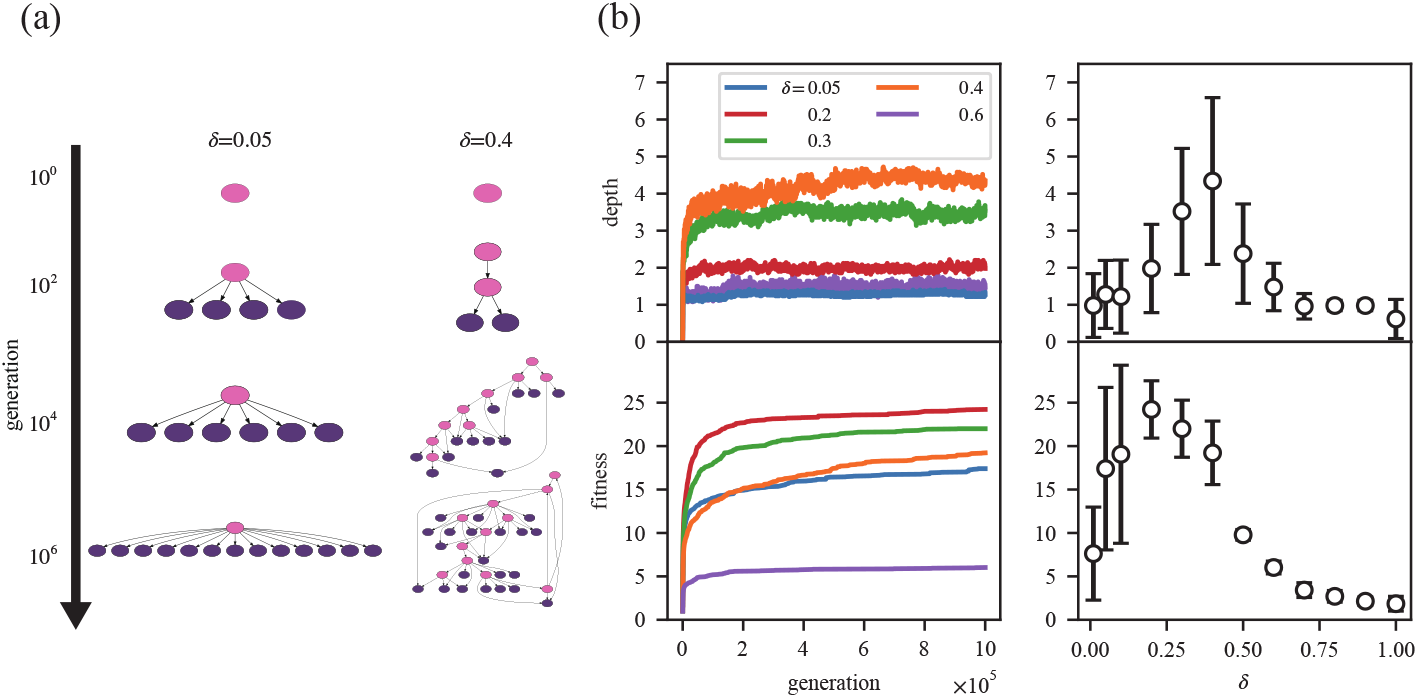
(a). Typical examples of time series of differentiation diagrams in GA under different *δ* (= 0.05, 0.4). (b)(left). The mean fitness and depth during GA. (right). The mean (white circle) and standard deviation (black bar) of fitness and depth reached at the final generation (= 10^6^). Each *δ* has 50 independent series. Depending on the value of *δ*, the depth and the fitness that emerge with optimization varies. Specifically, as *δ* changes, different types of diagrams emerged, including low-fitness and low-depth (*δ ≈* 0.01), high-fitness and low-depth (0.05 ⪅ *δ* ⪅ 0.2), high-fitness and high-depth (0.3 ⪅ *δ* ⪅ 0.4), low-fitness and low-depth (0.5 ⪅ *δ*). Note that all subsequent results in this paper are for *n* = 5 genes and *N* = 1000 perturbations.

Having discovered the clear trend of acquired depth along with the GA, the focus shifts to the reason why it appeared. In this model, cell differentiation occurs when perturbations cause the state to move out of the original attractor’s basin and into the adjacent attractor’s basin. While computing attractor basins is generally challenging, this model allows for a partial examination of basin structure the following two characteristics.

1. Since eq. (1) can be rewritten as follows, d***x***/d*t* of any point ***x*** takes the direction toward the lattice point ***a*** below.

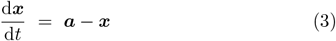

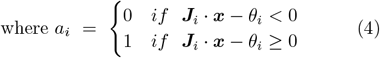

Property 1 is simply a substitution of ***a*** for the first term *H*(***J***_*i*_ ***x*** *−****θ***_*i*_) on the right-hand side in eq. (1). Here, in the two regions of the state space divided by the *i*-th hyperplane ***J***_*i*_ ***x*** *−θ*_*i*_ = 0, the *i*-th coordinate of the destination switches. Note that each of the *n* genes has the corresponding hyperplane. Fig. 3 illustrates this property using a two-dimensional example. When ***J***_*i*_ ***p*** *−θ*_*i*_ and ***J***_*i*_ ***q*** *−θ*_*i*_ have the same sign, ***p*** and ***q*** are referred to as being “on the same side” with respect to the hyperplane ***J***_*i*_ ***x*** *−θ*_*i*_ = 0.

The model then has the following property 2.

1. If the lattice point ***a*** is a fixed point and ***x*** is on the same side as ***a*** with respect to all *n* hyperplanes, then ***x*** converges to ***a*** after sufficient time.

**FIG. 3.**
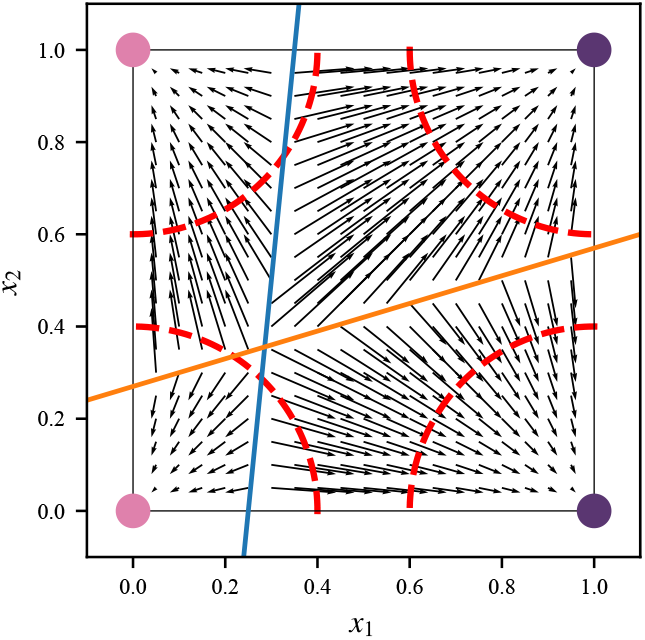
An example of state space in a two-gene system. d***x***/d*t* of the state space is visualized as a vector field. Two “hyperplanes”, ***J***_*i*_ *·* ***x*** *− θ*_*i*_ = 0 (*i* = 1, 2) are drawn as blue and orange lines respectively. The four points at each corner in blue are fixed point attractors. The sets of possible states that can be reached by perturbations are represented by red dashed lines that form an arc of radius *δ* in the 2D state space (in *n* dimensions, it is the surface of a hypersphere). When a state crosses the *i*-th hyperplane by being added some perturbation, the attractor to which it heads changes, and the *i*-th gene of the destination flips its value.

From Property 1, the time derivative of ***x*** is oriented toward ***a***, hence ***x*** moves toward ***a*** in a straight line. Since ***x*** and ***a*** are on the same side with respect to all of the hyperplanes, the line does not cross any of these hyperplanes on its way from ***x*** to ***a***.

As long as perturbations to an attractor do not cross any hyperplanes, the attractor will not transition to another attractor. In other words, if the hypersphere with a radius of *δ* centered on an attractor does not intersect any hyperplanes, it becomes a terminally differentiated state. Thus, increasing the number of terminally differentiated states requires that as many hyperspheres as possible do not intersect the hyperplane. In addition, points in the state space often converge to lattice points with coordinates of 0 or 1, since *H* in the eq. (1) is a step function^1^.

Using these properties, we can explain the difference in hierarchical structure between small and relatively large *δ* as follows. First, in order not to intersect as many hyperspheres as possible for the sake of increasing fitness under *δ* (*≈* 0.4), each hyperplane must be placed orthogonally to the corresponding axis, dividing the hypercube in two (Fig. 4a). When the planes are orthogonalized, the off-diagonal terms *J*_*ij*_(*j* ≠ *i*) are small relative to the diagonal term *J*_*ii*_. This can be understood from the fact that in the limit where *J*_*ij*_*/J*_*ii*_ *≪*1, the equation of the *i*-th hyperplane becomes *x*_*i*_ *− θ*_*i*_*/J*_*ii*_ = 0. Indeed, under the condition of *δ* = 0.4, the magnitude of the off-diagonal terms of *J* becomes much smaller than the diagonal terms (i.e., more orthogonal) as optimization proceeds (Fig. 4d). When the corresponding hyperplanes are orthogonalized with respect to each axis and separating adjacent hyperspheres, the intersection of these *n* hyper-planes will be located near the center of the hypercube. (the white dot in Fig. 4a). This idea is supported by the fact that, under the condition of *δ* = 0.4, the distribution of the distance between the intersection and ***a***^root^ tends to peak around 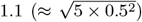 as optimization progresses (Fig. 4e). Since this is greater than *δ*(= 0.4), it becomes impossible to cross at least *n*(= 5) hyperplanes simultaneously. Moreover, the number of hyperplanes that can be crossed at once by a single perturbation becomes limited. As a result, in order to reach a terminally differentiated state that is distant from the root attractor, a minimum of two consecutive differentiation steps must be traversed. This phenomenon contributes partially to the reason behind the hierarchical nature of the multi-step differentiation diagram.

**FIG. 4.**
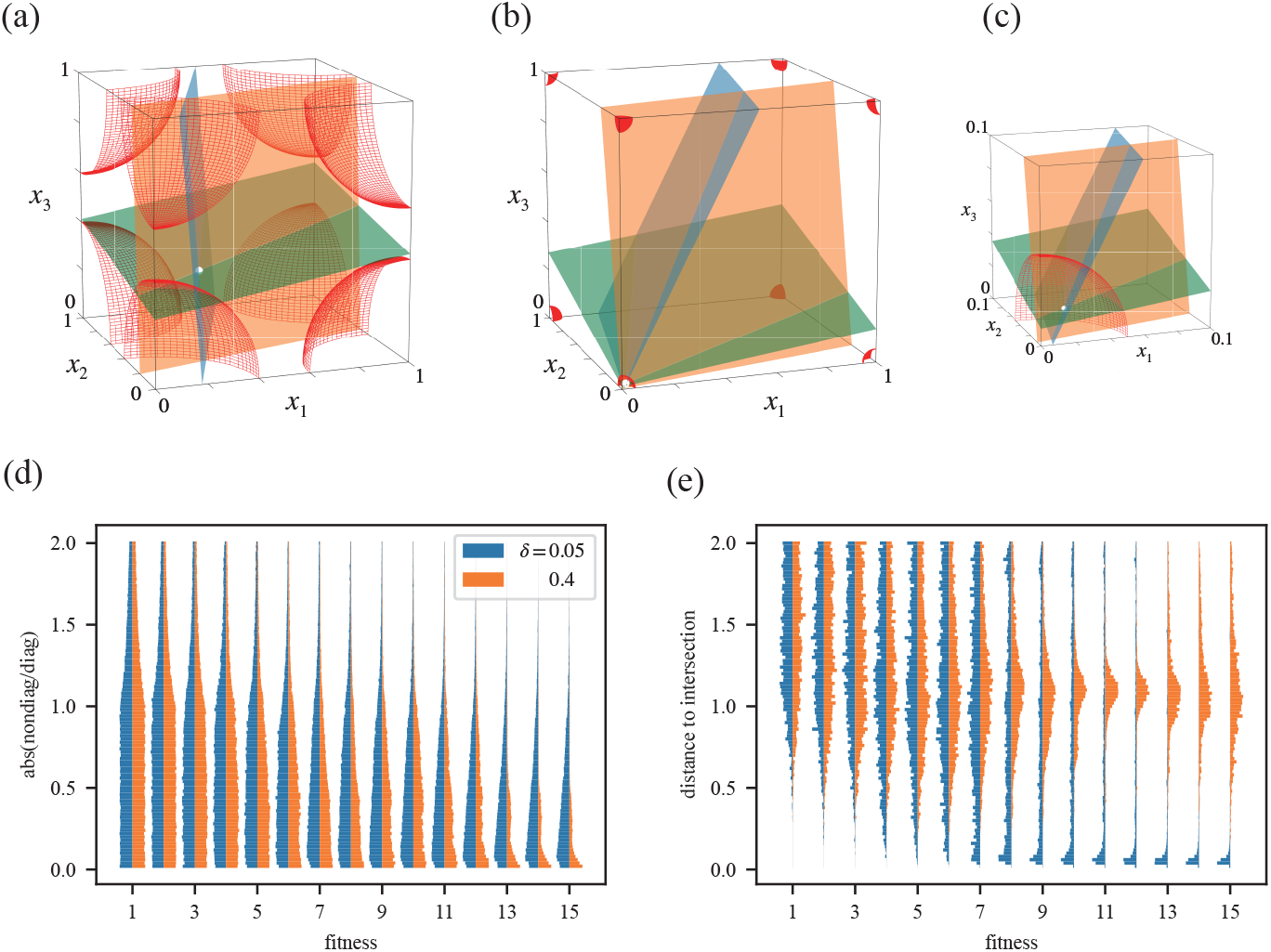
(a,b,c). Typical examples of 3D state spaces obtained from optimizing three-gene systems with *δ* = 0.4 (a) and 0.05 (b, c), respectively. Here, we use three-gene systems for visualization. Each plane represents ***J***_*i*_ *·* ***x*** *− θ*_*i*_ = 0 (*i* = 1 (blue), 2 (orange), 3 (green)). The white dot denotes the intersection of the three planes. The sets of possible states that can be reached by perturbations are represented by red octants. (c) is enlarged view near the origin in (b). In the case of *δ* = 0.4, the intersection is outside the sphere, whereas in the case of *δ* = 0.05, it is inside the sphere. (d) The distribution of the ratio *J*_*ij*_*/J*_*ii*_(*j≠ i*). The off-diagonal terms are low compared to the diagonal terms when it has high fitness under *δ* = 0.4, but not so much under *δ* = 0.05. (e) The distribution of the distance to the intersection from ***a***^root^ (, regardless of whether the intersection is inside the hypercube or not). Under *δ* = 0.4, samples with higher fitness tends to keep the distance farther away, while under *δ* = 0.05, the distance is optimized to be closer. In (d) and (e), we used multicanonical Monte Carlo method to sample 10000 systems for every fitness.

On the other hand, if *δ* is small, the hyperplanes can be placed more easily not intersecting the hyperspheres. This means that each hyperplane need not be orthogonalized during optimization (Fig. 4b,c,e). The intersection of the *n* hyperplanes can be placed close to the root attractor (Fig. 4b-d) since there is no such restriction for the hyperplanes to be orthogonalized as the case in *δ* = 0.4. In this case, it is unnecessary to create any precursor state to gain fitness. In addition, the hyperplanes are less likely to intersect a hypersphere of a relatively small radius. Thus, the non-hierarchical pattern of differentiation diagrams holds the majority.

### Acquisition of irreversibility

The second purpose of this work is to elucidate the condition and the mechanism by which irreversibility appears in differentiation diagrams.

The differentiation diagram in Fig. 2a seems to differentiate irreversibly. Hence, we will examine whether the high-fitness differentiation diagrams under *δ* = 0.4 actually have an irreversible tendency. Here, we focus on the graph only between precursor states since the terminally differentiated states do not differentiate further by definition and therefore cannot be included in a cycle (Fig. 5a). To quantify irreversibility of differentiation diagram, we use the size of the feedback arc set (FAS), which is the smallest set of edges that must be removed to make a graph acyclic. Using this measure, we show that high-fitness differentiation diagrams tend to have a smaller FAS when compared to randomly sampled graphs with the same number of precursor states (Fig. 5b). This trend is independent of the number of precursor states. In other words, even though there is no explicit selection pressure on irreversibility between precursor states, it inevitably arises due to the demand to increase the number of terminally differentiated states.

**FIG. 5.**
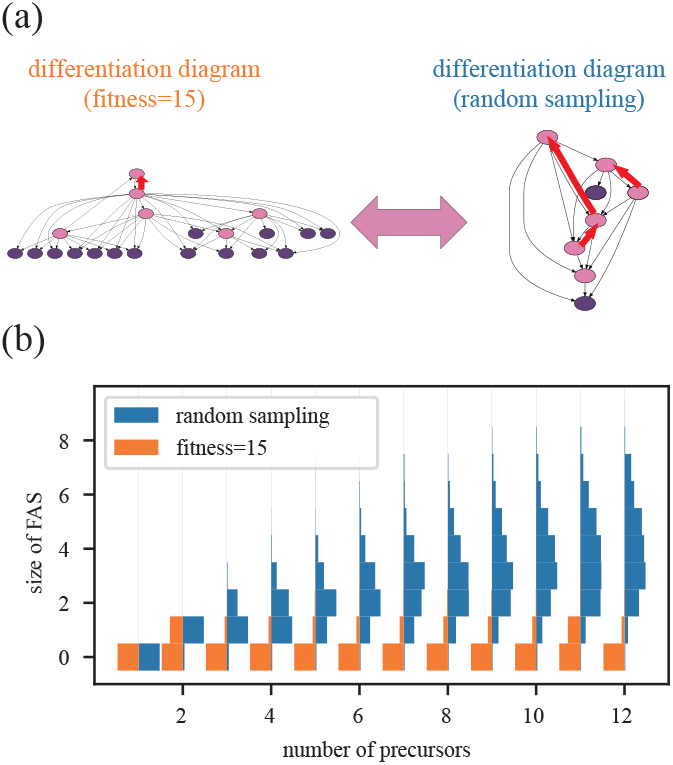
(a). A schematic of the comparison. We compared diagrams with fitness=15 to randomly sampled ones with the equivalent number of precursor states (drawn in pink). The red edges are an example of a feedback arc set (FAS) for each diagram. (b). The density of differentiation diagrams under *δ* = 0.4 with various numbers of precursor states (top) and their size distributions of FAS (bottom). Regardless of the number of precursor states, high-fitness (fitness=15) diagrams tend to have smaller FAS sizes compared to those of randomly sampled ones. Here, we used the multicanonical Monte Carlo method to collect 10000 samples in total for each.

What is the reason for the acquisition of irreversibility under *δ* = 0.4? In other words, why cannot diagrams have multiple cycles? In the following section, we will illustrate that the difficulty in creating cycles on a differentiation diagram can be accounted for by the orthogonalization of each hyperplane. Specifically, we focus particularly on the observation that in order to complete a cycle, i.e., to return to the original attractor by differentiation, the expression state of at least one gene must change in both the positive (0*→*1) and negative (1*→*0) directions.

As mentioned in the previous section, when the hyperplanes are orthogonalized with respect to the corresponding axes, the off-diagonal terms *J*_*ij*_(*j* ≠ *i*) are small relative to the diagonal term *J*_*ii*_. Hence, if the *i*-th plane is orthogonalized, the distance between an attractor ***a*** and the plane is approximated by the following equation.

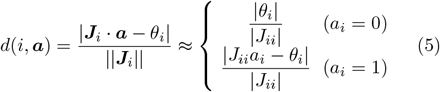

Here, this distance *d*(*i*, ***a***) is a typical threshold for the norm of a perturbation needed to differentiate from ***a*** to another attractor ***ā***^*i*^ whose *i*-th component is opposite to ***a***(Fig. 6c(inset)):

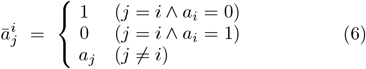

If *δ* is larger than *d*(*i*, ***a***), ***a*** can be differentiated into **ā**since a perturbation can cross the *i*-th hyperplane. Here, from approximation (5), *d*(*i*, ***a***) + *d*(*i*, **ā**) *≈*1 (Fig. 6c(inset)). This means that when *δ* ⪅ 0.5, differentiations in both directions (0 *→*1 and 1 *→*0) can hardly be possible at the same time. In addition, on the right-hand side of (5), the formula is independent of *a*_*j*_(*j* ≠ *i*). Hence, approximation (5) suggests that if the hyperplanes are orthogonalized, most of the differentiation paths which involve the *i*-th gene, regardless of *a*_*j*_, tend to head in the same direction (either 0 *→*1 or 1 *→*0) (Fig. 6a). Since the expression state of each gene of the root attractor 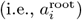 is likely to be zero^2^, differentiation from ***a***^root^ often occurs in the positive direction (0 *→*1), but not often in the negative direction (1 *→*0). This local tendency also appears globally in the state space due to the fact that the approximation of *d*(*i*, ***a***) is independent of *a*_*j*_(*j ≠ i*). In fact, in the optimized samples, more paths differentiate in the positive direction than in the negative direction (Fig. 6b). The distribution of the distances *d*_0_ from the attractor with *a*_*i*_ = 0 to each hyperplane and *d*_1_ from the one with *a*_*i*_ = 1 supports the hypothesis above (Fig. 6c). Thus, it is difficult to create a cycle because of such a bias in the direction of differentiation.

**FIG. 6.**
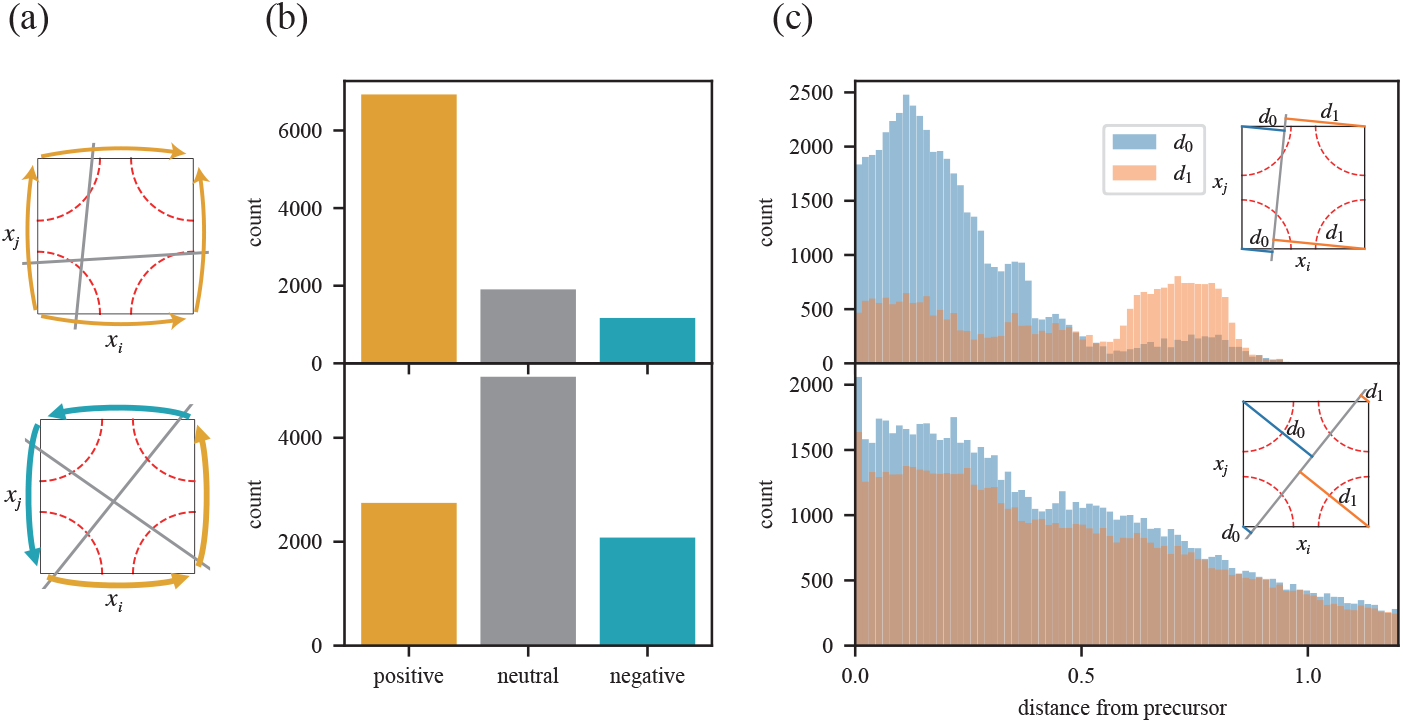
(a). An illustration of the mechanism how irreversibility emerges. Differentiations in positive direction are drawn in yellow, while negative are in cyan. The direction of differentiation in state space is biased positively with respect to each axis as the hyperplane is orthogonalized. (b). The bias of the differentiation direction in high-fitness samples (top) and randomly sampled ones (bottom). In high-fitness samples, the direction of differentiation is positively biased. On the other hand, in the random samples, there is no significant bias. (c). Distribution of *d*_0_ and *d*_1_ in high-fitness samples (top) and random samples (bottom). It indicates that in high-fitness samples, *d*_0_ has a peak in low value while *d*_1_ has a peak in high value. On the other hand, in random samples, there is no significant difference in the distribution of *d*_0_ and *d*_1_. The inset is a schematic visualization of *d*_0_ and *d*_1_. It shows that when the hyperplanes are orthogonalized, the distribution of *d*_0_ and *d*_1_ should be biased (if *d*_0_ is small, *d*_1_ should be relatively large). Here in (b,c), the high-fitness sample has fitness=15. We compared high-fitness and random samples with 5 precursor states, using 10000 samples for each.

### Structure of optimized

*GRNs* In the previous sections, we have seen that the key to the emergence of hierarchy and irreversibility is the diagonal terms’ (*J*_*ii*_) being much larger than the off-diagonal terms (*J*_*ij*_) in the interaction matrix. Next, can we find any rules here for the relationship between the off-diagonal terms? In the following, we will analyze the structure between the off-diagonal terms in more detail.

After optimization under *δ* = 0.05, no clear structure is observed in GRNs, while under *δ* = 0.4, there spontaneously appears a structure with a certain type of genes that receives interaction from other genes but gives little (less than a given threshold, 0.2) (Fig. 7a,b). We will refer to this type as “peripheral genes” (green in Fig. 7) and those that are not as “core genes” (gray in Fig. 7). The fact that peripheral genes receive little interaction means that the hyperplanes corresponding to the core genes are almost parallel to the axes of peripheral genes (Fig. 7c). In other words, the boundaries that determine the expression of core genes are almost independent of the expression states of peripheral genes. Hence, the two subspaces, one in which peripheral gene expression is on and the other in which is off, are almost identical (Fig. 7c,d). In short, peripheral genes play such a role as to clone the subspace spanned by core genes. Here, the peripheral genes divide the two subspaces in a way that makes it difficult for each state to differentiate in a reversible manner. In this way, when the state space is sliced in two by a peripheral gene, the original differentiation diagram can be copied in the direction of the peripheral gene (Fig. 7c,d). By introducing a hyperplane of a peripheral gene that intersects the center of the hypercube, with a slope of small but finite magnitude against the corresponding axis, it becomes feasible to enter the clonal subspace defined by the core genes through differentiation. This strategic approach holds the potential to ideally double the count of terminally differentiated states associated with the original differentiation diagram, without considering the peripheral gene. Importantly, this gene configuration can serve as a sufficient condition for constructing a hierarchical differentiation diagram capable of generating numerous terminally differentiated states.

**FIG. 7.**
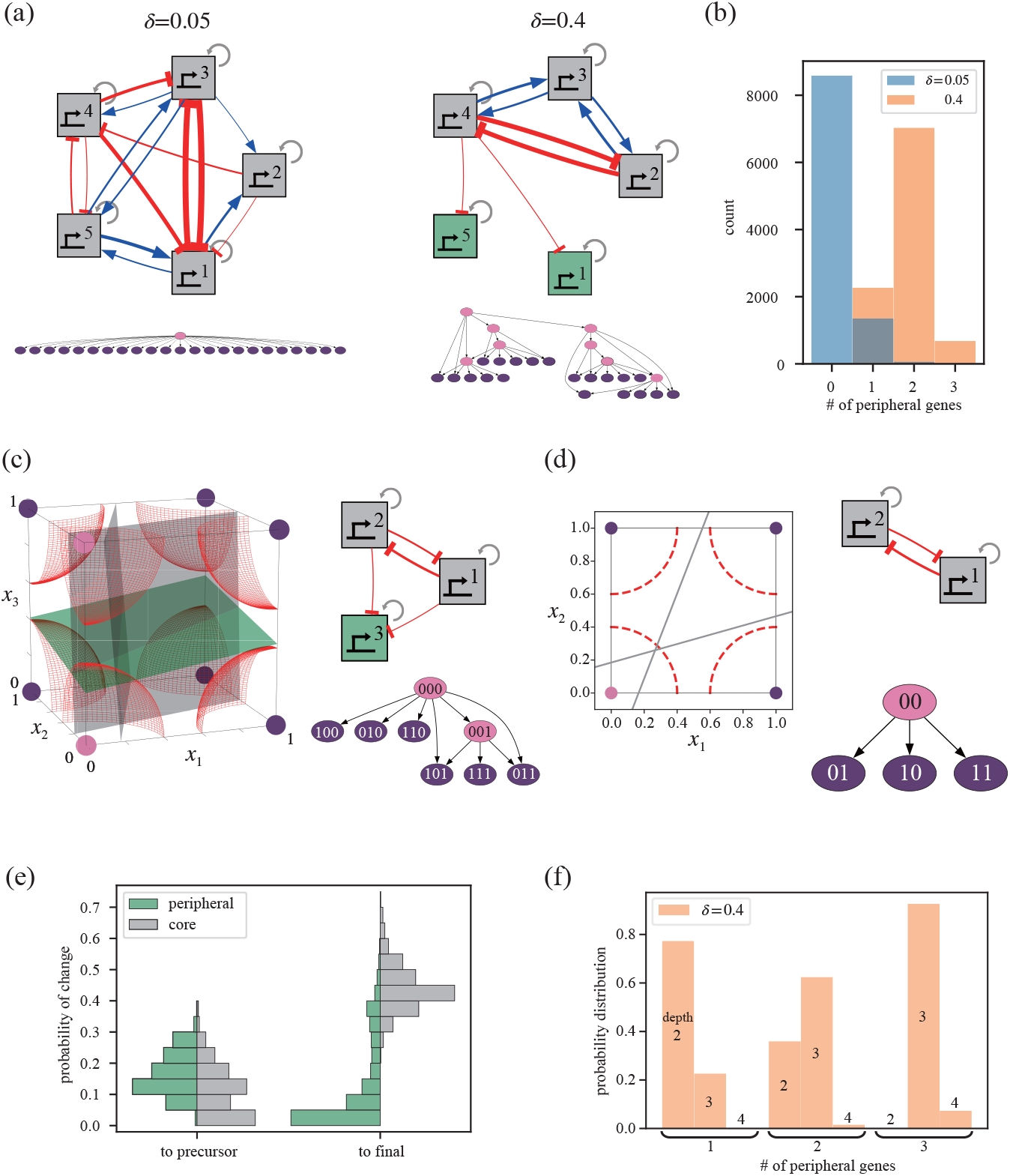
(a). Typical examples of GRNs (top) and the corresponding differentiation diagram (bottom) evolved under *δ* = 0.05, 0.4. Under *δ* = 0.4, in this example, there are two genes (gene 1 and 5, drawn in green) which interact little with any other gene. Here, auto-regulations (*J*_*ii*_) are drawn in gray arrows. Mutual regulations (*J*_*ij*_) are drawn in blue if positive and in red if negative. The widths of the arrows for mutual regulations are proportional to 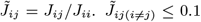 are omitted for visual clarity. (b). The distribution of the number of peripheral genes of networks with fitness = 20. Here, Peripheral genes are defined as those with an interaction value of 0.2 or less for any other gene (i.e., 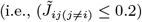 for all *j ≠ i*). (c).The difference between core and peripheral genes in a three-dimensional example. The colors of the planes correspond to those of the nodes in the GRN. The differentiation diagram of this example is shown in the lower right corner. (d). The state space, GRN, and differentiation diagram when the peripheral gene ‘3’ in (c) is excluded. By comparing it with (c), it can be observed that the state space formed by core genes is being replicated by the peripheral gene. (e). Distribution of the probability whether or not a single differentiation changes the expression state of each type of gene. In the case of differentiation to the precursor state, peripheral genes are slightly more likely to be involved. However, in the case to the terminally differentiated state, core genes are more likely to change, while peripheral genes change only rarely. (f). The correlation between depth of the diagram and the number of peripheral genes. Note that all samples presented here has fitness=20. We used multicanonical Monte Carlo method to collect 10000 samples for each *δ*.

Next, we investigate whether the described mechanism is utilized in the GA simulation. If this scenario is indeed employed, we would expect the peripheral genes to play a predominant role in differentiating between precursor states, while the core genes would primarily contribute to differentiation towards the terminally differentiated state. The reason behind this lies in the fact that peripheral genes play a key role in transitioning to the cloned subspace, but once core genes’ expression levels change, the traversal between subspaces along the axis of peripheral genes becomes difficult. This scenario finds support in the contrasting preferences observed regarding the types of genes more likely to be utilized in differentiation between precursor states and differentiation towards the final state (Fig. 7e).

In addition, the number of peripheral genes and the depth of the diagram has a positive correlation (Fig. 7f). This is because the more peripheral genes there are, the more differentiation there is using those peripheral genes before using core genes to differentiate into the terminally differentiated state. To increase the number of terminally differentiated states, peripheral genes have such an effect that the system of the core network is used repeatedly.

The “core & peripheral” strategy is used in the majority of cases. This fact suggests that the parameter sets in such a strategy exists much more than those which use all the genes as core.

## DISCUSSION

In this paper, we employed an abstract cell model to study the characteristics which arise as the differentiation diagrams evolve. Here, the cell state is regulated by GRNs, and differentiation is induced by perturbation of the gene expression, whose strength is represented as *δ*. We revealed that the emergence of hierarchy and irreversibility in differentiation diagrams depends on *δ*. Our findings are novel in that they demonstrate the inevitability of these properties through optimization of the number of terminally differentiated states. Specifically, under conditions of high gene expression noise, these properties will inevitably emerge. In fact, several researches reported that the magnitude of stochastic fluctuations in gene expression in living cells can reach levels comparable to the average expression level [35, 36]. Thus, actual organisms might have evolved in an condition with such large fluctuations in expression. The outcomes of evolution under these highly fluctuating conditions may be effectively captured by our theoretical model.

Next, we further elucidated the mechanism behind the acquisition of hierarchy and irreversibility in differentiation diagrams. To investigate the mechanism, we presented a geometric approach in high-dimensional state space. Specifically, we considered the intersection of the hyperspheres centered on each attractor and the hyperplanes determined by the GRN. This helped us to determine whether perturbations could cross typical basin boundaries and cause a differentiation. The results showed that when fluctuations were high, the hyperplanes were orthogonal to the axis for attractors not to cross the basin boundary by perturbations. This orthogonalization means that each gene generally had stronger positive self-regulation compared to the mutual regulation with other genes. Indeed, several studies on key genes that regulate cell differentiation have indicated that the estimated strength of positive self-regulation exceeds the strength of mutual regulation from other genes [37, 38]. However, there is still a limited number of quantitative assessments on gene interactions in the context of GRN inference, requiring further extensive examination.

We also observed that a “core & peripheral” structure emerged in the GRN after optimization under *δ* = 0.4. Here, the genes were spontaneously divided into two groups, referred to as core genes and peripheral genes, breaking the initial symmetry between genes. The system was typically structured to change the expression of peripheral genes while transitioning between precursors, while the state of core genes was changed mainly in transition into a terminally differentiated state. This system uses the same genes in various transition events, and distinguishes cell types through different combinations of gene expression states. In fact, recent data from vertebrate development partially supports this idea of generating various stable cell types through combinatorial reuse of transcription factors [4]. On the other hand, the process of cell fate determination has been majorly explained by the expression of lineage-specific genes which are regulated by bistable switches [39–41]. These switches control the expression of genes, and their hierarchical linkages are thought to lead the multi-step process of cell fate determination [25, 42]. However, recent researches suggested that the inferred structure of GRN controlling hematopoiesis is somehow densely interconnected rather than hierarchical [38, 43]. Here, Our model can provide a fresh perspective and prediction on the GRN structure that generates hierarchical differentiation. Further research will be required to verify whether this novel mechanism can be found in actual organisms.

Considering the intrinsic stochastic nature of gene ex-pression in actual organisms [44, 45], this study employed a model that considers fluctuations in gene expression as the primary driving force behind transitions between stable states. In fact, this concept does not contradict the evidence from various studies that have demonstrated the role of stochastic gene expression in hematopoietic differentiation [46–48] and the early developmental process of mice [49, 50]. Since our model is not specific to any particular genes, signals, or environmental conditions, its conclusions might be applied to a broad range of systems where stochastic state transitions can occur.

## MATERIALS & METHODS

### Genetic algorithm

We executed a genetic algorithm with a population size of 100. In each generation, the top 20 individuals were selected. From each of these 20 individuals, four mutated offspring were generated, resulting in a total of 100 individuals for the next generation, including the original parents. The mutation process involved introducing new values to each element of (*J*, ***θ***) with a probability of 0.05. The new values were sampled from a uniform distribution in the range of [*−*1, 1].

### Multicanonical Markov chain Monte Carlo method

This approach facilitates the effective sampling of rare events, including low-energy states, while avoiding entrapment in local minima. For both advantages of an optimization algorithm (energy minimization) and unbiased random sampling (energy distribution calculation), this method has been applied beyond the field of physics [34, 51, 52]. Notably, this method has also found utility in the context of the evolution of GRN [53– 55]. In our study, we consider the Markov chain sampling process for (*J*, ***θ***) states, represented as (*J*, ***θ***) *→* (*J* ^*′*^, ***θ***^*′*^) *→*…. Here, the new candidate state (*J* ^*′*^, ***θ***^*′*^) is generated by mutating elements of the former state (*J*, ***θ***), based on the same algorithm used in the GA. Specifically, we assigned a mutation probability of 0.05 for each element of (*J*, ***θ***), with new values sampled from a uniform distribution in the range of [−1, 1]. The transition from (*J*, ***θ***) to (*J*, ***θ***^*′*^) is determined by the transition probability function Π_*f*(*J*,***θ***)*→f*(*J*′,***θ***′)_ = min (1, *w*(*f* (*J*′, ***θ*** ′))*/w*(*f* (*J*, ***θ***))), which decides whether to accept or reject the candidate state based on its fitness-dependent weight function *w*. If an sufficient amount of samples has been collected, the sampled fitness distribution *P*_*eq*_(*f* (*J*, ***θ***)) follows the detailed balance condition Π_*f→f*_′ *P*_*eq*_(*f* (*J*, ***θ***)) = Π_*f*_′*→f P*_*eq*_(*f* ′ (*J* ′, ***θ*** ′)). Apparently, this distribution *P*_*eq*_(*f*) depends on the weight function *w*. In multicanonical sampling, the multicanonical weight *w*(*f*) *∝*1*/*Ω(*f*) is used. This leads to a uniform marginal distribution *h*(*f*) *∝w*(*f*) Ω(*f*) = const., which indicates a sampled histogram of the fitness approaches asymptotically to a uniform distribution. Hence, within the framework of multicanonical sampling, the Markov chain-generated sequence of (*J*, ***θ***) can be regarded as a random walk in the fitness space, enabling the efficient sampling of rare events (i.e., high-fitness states) without becoming trapped in local minima.

To calculate the multicanonical weight required for obtaining a flat histogram with respect to fitness, we employ the Wang-Landau algorithm [56] in advance. In this algorithm, it is necessary to predefine and discretize the range of the cost function *f* . In this study, we set the range to [1, 20] with a bin width of 1. Furthermore, this algorithm includes an operation to assess the “flatness” of the obtained fitness histograms. Here, as the criterion for flatness, we considered the histogram to be “sufficiently flat” when the counts of all bins exceed 90% of the expected value for a perfectly flat histogram.

## ACKNOWLEDGMENTS

This work was supported by RIKEN Junior Research Associate Program (to YTN), the Japan Society for Promotion of Science (JSPS) KAKENHI (17H06389, 22K21344 to CF; 22K15069, 22H05403 to YH), Japan Science and Technology Agency (JST) ERATO (JPM-JER1902 to CF), and Cooperative Study Program of Exploratory Research Center on Life and Living Systems (ExCELLS; program No. 20-102, 21-102 to NS). We thank Y. Uchida, S. Tsuru and T. Kohsokabe for stimulating discussion.

## Footnotes

1 Despite *H* being a step function, it is possible to have exceptional cases where it does not converge to a lattice point. An example of such a case is when a vector field is formed in a direction facing across a hyperplane. However, these cases become less frequent with optimization in this study. For instance, when the fitness is 15, such cases exist in less than 1% of cases for both *δ* = 0.05, 0.4.

2 For instance, under *δ* = 0.4 and fitness being 15, the percentage of 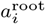 that are not 0 is less than 0.01%.

## Notes

### Competing Interest Statement

The authors have declared no competing interest.

## Reference

[1] L. Wolpert, C. Tickle, and A. M. Arias, Principles of development (Oxford University Press, USA, 2019) pp. 333–396.

[2] B. K. Tusi, S. L. Wolock, C. Weinreb, Y. Hwang, D. Hidalgo, R. Zilionis, A. Waisman, J. R. Huh, A. M. Klein, and M. Socolovsky, Population snapshots predict early haematopoietic and erythroid hierarchies, Nature 555, 54 (2018).

[3] D. E. Wagner, C. Weinreb, Z. M. Collins, J. A. Briggs, S. G. Megason, and A. M. Klein, Single-cell mapping of gene expression landscapes and lineage in the zebrafish embryo, Science 360, 981 (2018), https://www.science.org/doi/pdf/10.1126/science.aar4362.

[4] J. A. Briggs, C. Weinreb, D. E. Wagner, S. Megason, L. Peshkin, M. W. Kirschner, and A. M. Klein, The dynamics of gene expression in vertebrate embryogenesis at single-cell resolution, Science 360, eaar5780 (2018), https://www.science.org/doi/pdf/10.1126/science.aar5780.

[5] M. Plass, J. Solana, F. A. Wolf, S. Ayoub, A. Misios, P. Glaž ar, B. Obermayer, F. J. Theis, C. Kocks, and N. Rajewsky, Cell type atlas and lineage tree of a whole complex animal by single-cell transcriptomics, Science 360, eaaq1723 (2018), https://www.science.org/doi/pdf/10.1126/science.aaq1723.

[6] J. A. Farrell, Y. Wang, S. J. Riesenfeld, K. Shekhar Regev, and A. F. Schier, Single-cell reconstruction of developmental trajectories during zebrafish embryogenesis, Science 360, eaar3131 (2018), https://www.science.org/doi/pdf/10.1126/science.aar3131.

[7] J. S. Packer, Q. Zhu, C. Huynh, P. Sivaramakrishnan, E. Preston, H. Dueck, D. Stefanik, K. Tan, C. Trapnell, J. Kim, R. H. Waterston, and J. I. Murray, A lineage-resolved molecular atlas of ¡i¿c. elegans¡/i¿ embryogenesis at single-cell resolution, Science 365, eaax1971 (2019), https://www.science.org/doi/pdf/10.1126/science.aax1971.

[8] S. Siebert, J. A. Farrell, J. F. Cazet, Y. Abeykoon, S. Primack, C. E. Schnitzler, and C. E. Juliano, Stem cell differentiation trajectories in ¡i¿hydra¡/i¿ resolved at single-cell resolution, Science 365, eaav9314 (2019), https://www.science.org/doi/pdf/10.1126/science.aav9314.

[9] K. Shekhar, I. E. Whitney, S. Butrus, Y.-R. Peng, and J. R. Sanes, Diversification of multipotential postmitotic mouse retinal ganglion cell precursors into discrete types, eLife 11, 10.7554/eLife.73809 (2022).

[10] R. Shahan, C.-W. Hsu, T. M. Nolan, B. J. Cole, I. W. Taylor, L. Greenstreet, S. Zhang, A. Afanassiev, A. H. C. Vlot, G. Schiebinger, P. N. Benfey, and U. Ohler, A single-cell arabidopsis root atlas reveals developmental trajectories in wild-type and cell identity mutants, Developmental Cell 57, 543 (2022).

[11] C. H. Waddington, The strategy of the genes. A discussion of some aspects of theoretical biology. With an appendix by H. Kacser. (London: George Allen & Unwin, 1957) pp. 11–58.

[12] K. Takahashi and S. Yamanaka, Induction of pluripotent stem cells from mouse embryonic and adult fibroblast cultures by defined factors, Cell 126, 663 (2006).

[13] S. Huang, I. Ernberg, and S. Kauffman, Cancer attractors: A systems view of tumors from a gene network dynamics and developmental perspective, Seminars in Cell & Developmental Biology 20, 869 (2009).

[14] Q. Li, A. Wennborg, E. Aurell, E. Dekel, J.-Z. Zou, Y. Xu, S. Huang, and I. Ernberg, Dynamics inside the cancer cell attractor reveal cell heterogeneity, limits of stability, and escape, Proceedings of the National Academy of Sciences 113, 2672 (2016).

[15] E. H. Davidson, Gene Regulatory Networks and the Evolution of Animal Body Plans, Science 311, 796 (2006).

[16] G. Swiers, R. Patient, and M. Loose, Genetic regulatory networks programming hematopoietic stem cells and erythroid lineage specification, Developmental Biology 294, 525 (2006).

[17] K. Kobayashi, K. Maeda, M. Tokuoka, A. Mochizuki, and Y. Satou, Controlling cell fate specification system by key genes determined from network structure, iScience 4, 281 (2018).

[18] S. A. Kauffman, Metabolic stability and epigenesis in randomly constructed genetic nets, Journal of Theoretical Biology 22, 437 (1969).

[19] A. S. Ribeiro and S. A. Kauffman, Noisy attractors and ergodic sets in models of gene regulatory networks, Journal of Theoretical Biology 247, 743 (2007).

[20] I. Shmulevich, S. A. Kauffman, and M. Aldana, Eukaryotic cells are dynamically ordered or critical but not chaotic, Proceedings of the National Academy of Sciences 102, 13439 (2005), https://www.pnas.org/content/102/38/13439.full.pdf.

[21] R. Serra, M. Villani, A. Graudenzi, and S. Kauffman, Why a simple model of genetic regulatory networks describes the distribution of avalanches in gene expression data, Journal of Theoretical Biology 246, 449 (2007).

[22] S. Huang, Y.-P. Guo, G. May, and T. Enver, Bifurcation dynamics in lineage-commitment in bipotent progenitor cells, Developmental Biology 305, 695 (2007).

[23] S. Huang, Reprogramming cell fates: reconciling rarity with robustness, BioEssays 31, 546 (2009).

[24] O. Cinquin and J. Demongeot, Positive and negative feedback: Striking a balance between necessary antagonists, Journal of Theoretical Biology 216, 229 (2002).

[25] T. G. W. Graham, S. M. A. Tabei, A. R. Dinner, and I. Rebay, Modeling bistable cell-fate choices in the Drosophila eye: qualitative and quantitative perspectives, Development 137, 2265 (2010).

[26] M. N. Artyomov, A. Meissner, and A. K. Chakraborty, A model for genetic and epigenetic regulatory networks identifies rare pathways for transcription factor induced pluripotency, PLOS Computational Biology 6, e1000785 (2010).

[27] J. Krumsiek, C. Marr, T. Schroeder, and F. J. Theis, Hierarchical differentiation of myeloid progenitors is encoded in the transcription factor network, PLOS ONE 6, e22649 (2011).

[28] C. Furusawa and K. Kaneko, Robust development as a consequence of generated positional information, Journal of Theoretical Biology 224, 10.1016/S0022-5193(03)00189-9 (2003).

[29] N. Suzuki, C. Furusawa, and K. Kaneko, Oscillatory protein expression dynamics endows stem cells with robust differentiation potential, PLOS one 6, e27232 (2011).

[30] Y. Matsushita and K. Kaneko, Homeorhesis in Waddington’s landscape by epigenetic feedback regulation, Physical Review Research 2, 023083 (2020).

[31] R. Zhu, J. M. del Rio-Salgado, J. Garcia-Ojalvo, and M. B. Elowitz, Synthetic multistability in mammalian cells, Science 375, 10.1126/science.abg9765 (2022).

[32] S. Huang, Non-genetic heterogeneity of cells in development: more than just noise, Development 136, 3853 (2009).

[33] L. A. Buttitta and B. A. Edgar, Mechanisms controlling cell cycle exit upon terminal differentiation, Current Opinion in Cell Biology 19, 697 (2007).

[34] Y. Iba, N. Saito, and A. Kitajima, Multicanonical mcmc for sampling rare events: an illustrative review, Annals of the Institute of Statistical Mathematics 66, 611 (2014).

[35] M. B. Elowitz, A. J. Levine, E. D. Siggia, and P. S. Swain, Stochastic gene expression in a single cell, Science 297, 1183 (2002), https://www.science.org/doi/pdf/10.1126/science.1070919.

[36] W. R. Holmes, N. S. Reyes de Mochel, Q. Wang, H. Du, T. Peng, M. Chiang, O. Cinquin, K. Cho, and Q. Nie, Gene expression noise enhances robust organization of the early mammalian blastocyst, PLOS Computational Biology 13, 1 (2017).

[37] S. Jang, S. Choubey, L. Furchtgott, L.-N. Zou, A. Doyle, V. Menon, E. B. Loew, A.-R. Krostag, R. A. Martinez, L. Madisen, B. P. Levi, and S. Ramanathan, Dynamics of embryonic stem cell differentiation inferred from single-cell transcriptomics show a series of transitions through discrete cell states, eLife 6, e20487 (2017).

[38] J. E. Handzlik and Manu, Data-driven modeling predicts gene regulatory network dynamics during the differentiation of multipotential hematopoietic progenitors, PLOS Computational Biology 18, 1 (2022).

[39] S. H. Orkin, Diversification of haematopoietic stem cells to specific lineages, Nature Reviews Genetics 1, 57 (2000).

[40] P. Laslo, C. J. Spooner, A. Warmflash, D. W. Lancki, H.-J. Lee, R. Sciammas, B. N. Gantner, A. R. Dinner, and H. Singh, Multilineage transcriptional priming and determination of alternate hematopoietic cell fates, Cell 126, 755 (2006).

[41] L. Wang, B. L. Walker, S. Iannaccone, D. Bhatt, P. J. Kennedy, and W. T. Tse, Bistable switches control memory and plasticity in cellular differentiation, Proceedings of the National Academy of Sciences 106, 6638 (2009), https://www.pnas.org/doi/pdf/10.1073/pnas.0806137106.

[42] J. X. Zhou, L. Brusch, and S. Huang, Predicting Pancreas Cell Fate Decisions and Reprogramming with a Hierarchical Multi-Attractor Model, PLOS ONE 6, 10.1371/journal.pone.0014752 (2011).

[43] N. Novershtern, A. Subramanian, L. N. Lawton, R. H. Mak, W. N. Haining, M. E. McConkey, N. Habib, N. Yosef, C. Y. Chang, T. Shay, G. M. Frampton, C. Drake, I. Leskov, B. Nilsson, F. Preffer, D. Dombkowski, J. W. Evans, T. Liefeld, J. S. Smutko, J. Chen, N. Friedman, R. A. Young, T. R. Golub, A. Regev, and L. Ebert, Densely interconnected transcriptional circuits control cell states in human hematopoiesis, Cell 144, 296 (2011).

[44] M. Thattai and A. van Oudenaarden, Intrinsic noise in gene regulatory networks, Proceedings of the National Academy of Sciences 98, 8614 (2001), https://www.pnas.org/doi/pdf/10.1073/pnas.151588598.

[45] E. M. Ozbudak, M. Thattai, I. Kurtser, A. D. Grossman, and A. van Oudenaarden, Regulation of noise in the expression of a single gene, Nature Genetics 31, 69 (2002).

[46] D. A. Hume, Probability in transcriptional regulation and its implications for leukocyte differentiation and inducible gene expression, Blood 96, 2323 (2000).

[47] H. H. Chang, M. Hemberg, M. Barahona, D. E. Ingber, and S. Huang, Transcriptome-wide noise controls lineage choice in mammalian progenitor cells, Nature 453, 544 (2008).

[48] J. C. Wheat, Y. Sella, M. Willcockson, A. I. Skoultchi Bergman, R. H. Singer, and U. Steidl, Single-molecule imaging of transcription dynamics in somatic stem cells, Nature 583, 431 (2020).

[49] J.-E. Dietrich and T. Hiiragi, Stochastic patterning in the mouse pre-implantation embryo, Development 134, 4219 (2007).

[50] A. Eldar and M. B. Elowitz, Functional roles for noise in genetic circuits, Nature 467, 167 (2010).

[51] N. Saito, Y. Iba, and K. Hukushima, Multicanonical sampling of rare events in random matrices, Phys. Rev. E 82, 031142 (2010).

[52] A. Kitajima and M. Kikuchi, Numerous but rare: An exploration of magic squares, PLOS ONE 10, 1 (2015).

[53] N. Saito and M. Kikuchi, Robustness leads close to the edge of chaos in coupled map networks: toward the understanding of biological networks, New Journal of Physics 15, 053037 (2013).

[54] S. Nagata and M. Kikuchi, Emergence of cooperative bistability and robustness of gene regulatory networks, PLOS Computational Biology 16, 1 (2020).

[55] T. Kaneko and M. Kikuchi, Evolution enhances mutational robustness and suppresses the emergence of a new phenotype: A new computational approach for studying evolution, PLOS Computational Biology 18, 1 (2022).

[56] F. Wang and D. P. Landau, Efficient, multiple-range random walk algorithm to calculate the density of states, Phys. Rev. Lett. 86, 2050 (2001).

